# The CAGE complex: a hollow, megadalton, protein assembly in prokaryotic and eukaryotic microbes

**DOI:** 10.1101/2025.09.22.677704

**Authors:** Caitlyn L. McCafferty, Gabriel Hoogerbrugge, Ophelia Papoulas, Evan A. Schwartz, Simone Ritchey, David W. Taylor, Axel F. Brilot, Edward M. Marcotte

## Abstract

We report the discovery and structure of a previously unknown ∼1 MDa hollow protein assembly, identified during a survey of ciliary complexes from the ciliate *Tetrahymena thermophila*. By combining mass spectrometry, structure prediction, and cryo-electron microscopy, we define a homotetrameric cage-like complex with a distinctive elliptical architecture and a large internal cavity. A sequence survey revealed several thousand homologs spanning diverse unicellular eukaryotes—including green algae, fungi, amoebozoans, choanoflagellates, and SAR lineages—as well as predominantly gram-negative bacteria, indicating an ancient evolutionary origin and arguing against a eukaryote-specific function. We determined a near-atomic resolution structure of the complex from the slime mold *Dictyostelium discoideum*, demonstrating conservation of overall architecture and cavity despite low sequence identity. Together, these results establish the CAGE complex (<u>C</u>onserved <u>A</u>ssembly in <u>G</u>ram-negative bacteria and <u>E</u>ukaryotes) as a new class of large protein cage broadly distributed across the tree of life. While its biological function remains unknown, its size, architecture, and conservation suggest possible roles in transport or protein/RNA homeostasis.

## Introduction

Microscopists have long relied on visual assessments of cell and tissue samples, and cryo-electron microscopy (cryo-EM) extends this capability to atomic resolution, offering unprecedented insights into molecular structures. Traditionally, structural biology studies are hypothesis-driven where there is focus on the purification of a single protein complex of interest. Alternatively, visual examinations of cells and cell lysates have historically led to the discovery of many particularly striking molecular assemblies, such as the giant polyhedral prokaryotic protein-based organelles known as carboxysomes^1^, the snake-like nucleotide biosynthesis enzyme fibers known as cytoophidia^2^, and the remarkable, 8 MDa, 39-fold symmetric vault complex found in many eukaryotic cells and so-named for its visual similarity to arched cathedral ceilings^3,4^.

Importantly, these sorts of surveys can generally be applied to any species or tissue, including less-well studied organisms^5–12^. One such study in our lab centered on protein machinery found in the swimming unicellular microbial eukaryote *Tetrahymena thermophila*, with a particular focus on determining proteins localized to the *Tetrahymena* cilia, the microtubule-based eukaryotic flagellum that provides motility to these microbes and is broadly conserved across eukaryotic organisms including humans^13^. In particular, we sought to focus our efforts on the most easily visualized macromolecular complexes by first biochemically separating native protein extracts using chromatography, then using cryo-electron microscopy (cryo-EM) to survey the biochemical fractions containing megadalton-scale assemblies^14^. These crude fractions were still highly heterogeneous in nature, generally containing tens to hundreds of protein complexes, as estimated using protein mass spectrometry. However, in doing so, we observed a striking macromolecular assembly within these extracts that did not obviously resemble any structures known to date, and we set out to characterize it.

In this paper, we describe the serendipitous discovery and structural determination of this novel cage-like multiprotein assembly, termed the CAGE complex. Subsequent sequence analysis and mass spectrometry investigations revealed its component proteins, which we named the CAGE family of proteins, to be widely distributed across the tree of life in both prokaryotic and eukaryotic, predominantly unicellular, microbes, including select fungi, green algae, the slime mold *Dictyostelium discoideum,* and the choanoflagellate *Salpingoeca rosetta*.

In order to confirm that the 3D architecture of CAGE proteins is conserved, we determined the high-resolution structure of the native CAGE complex from fractionated *Dictyostelium* cell lysate. Based on cross-linking mass spectrometry (XL/MS) observations and structural considerations, we hypothesize possible functions for the CAGE complex. This work underscores the significance of discovery-based biological surveys for broadening our understanding of complex cellular biology and points to many new structural assemblies still to be characterized across the tree of life.

## Results

### An electron microscopy survey reveals a hollow, cage-like protein assembly in the Tetrahymena ciliary matrix

Many structural studies of cilia are focused on the intricate microtubule-base ultrastructure and auxiliary molecular components, however, we were interested in the less-studied ciliary matrix. The cilia proteome is estimated to contain over 1,000 proteins^15,16^, with approximately 300 involved in the ciliary ultrastructure^17,18^, including axonemal microtubules, dynein arms, radial spokes, and other associated linkers. This leaves hundreds of proteins that are likely to localize to the ciliary membrane and matrix. While cryo-ET has proven an effective means of studying ciliary ultrastructure^19–21^, the ciliary matrix remains a more challenging environment for cellular tomography due to the lack of periodic complexes when compared to the ciliary axoneme. To better characterize matrix-associated proteins, we first isolated cilia from the multiciliated protist *Tetrahymena*, and then subfractionated the cilia into axoneme and membrane & matrix fractions (M&M) (**Figure 1A**). We then separated protein complexes in the M&M fraction using size exclusion chromatography (SEC) to further reduce the sample complexity before freezing selected fractions containing megadalton-sized complexes on cryo-EM grids (**Figure 1B-C**). We identified the most abundant proteins in the fractions using mass spectrometry (**Figure 1D**) and, in parallel, performed cross-linking mass spectrometry on the selected fractions to determine interactions and structural constraints among the proteins present.

**Figure 1.**
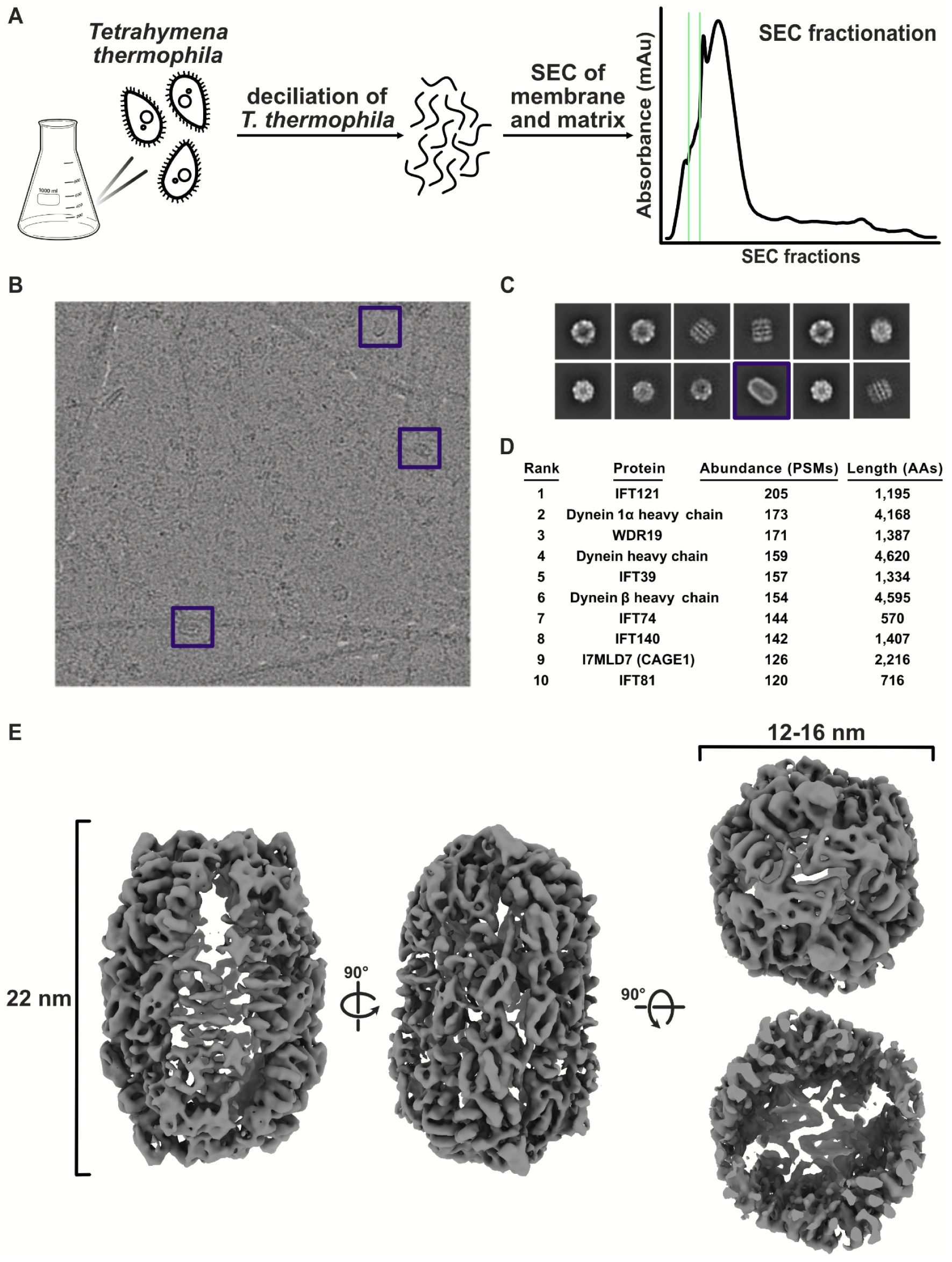
Cryo-EM of the ciliary matrix reveals a large, previously uncharacterized, hollow protein complex. **(A)** Cilia from *Tetrahymena thermophila* were isolated using dibucaine and further separated into axonemal, and membrane and matrix fractions. The membrane and matrix fraction was further simplified using size exclusion chromatography (SEC) from which the fractions containing large complexes were selected for imaging with cryo-EM. **(B)** Cryo-EM survey of the high molecular weight complexes shows diverse molecular species, including a large elliptical particle (purple box). **(C)** 2D classification of particles picked with a blob picker yields different views of a variety of complexes, with an example elliptical particle class again shown with a purple box. **(D)** Mass spectrometry of the imaged fractions, indicating the 10 most abundant proteins. **(E)** 3D reconstruction of the elliptical particle shown in **(B)** and **(C)** at 5.64 Å resolution reveals a large hollow protein shell with a long axis of approximately 22 nm and a diameter of 12-16 nm.

Particles were picked from the micrographs using a generalized blob picking approach^22^, which revealed several distinct molecular species upon 2D classification. These 2D classes revealed familiar complexes such as dynein heavy chains and the CCT chaperone, in addition to an unfamiliar elliptical complex measuring about 22 nm x 16 nm x 12 nm wide (**Figure 1C**). Particles assigned to this latter class were selected, the data reprocessed with a template created from the initial particles, and reconstructed into a D2 symmetric 3D map that was resolved to 5.64 Å resolution (**Figure 1E**).

### Identification of proteins comprising the CAGE complex

We examined the mass spectrometry-identified proteins in the biochemical fractions surveyed by electron microscopy to find potential candidates for the protein(s) composing the CAGE assembly (**Figures 1D, 2A; Supp Data**). These fractions were dominated by known ciliary machinery, primarily the Intraflagellar Transport (IFT) A and B complexes, as well as numerous dynein heavy chains (e.g. see ^23^) and chaperone complexes. However, among the 10 most abundant proteins in these fractions was a large, uncharacterized protein known only from its Uniprot database identifier, I7MLD7_TETTS. Given this lack of functional and structural characterization, we therefore next attempted to assess the candidate by computationally predicting its 3D structure.

Prediction of the structure of I7MLD7_TETTS using AlphaFold^24^ suggested that it was highly structured and might adopt a roughly quarter-spherical, cup-shaped fold with spatial dimensions broadly consistent with those observed for the hollow cage observed in our EM datasets (**Figure 2B**). As I7MLD7_TETTS eluted chromatographically above its predicted monomeric molecular weight of 258 kD, we examined the likelihood of a homomeric assembly by calibrating our size exclusion chromatography with molecular weight standards in order to experimentally estimate the molecular weight of I7MLD7_TETTS based upon its chromatographic elution position. The experimentally observed molecular weight of I7MLD7_TETTS was consistent with a multimeric assembly of 3-5 copies (**Figure 2C**). We further tested the possibility of homo-oligomerization by repeating structure prediction with AlphaFold Multimer^25^ and evaluating alternative stoichiometries. These tests strongly supported the possibility of a homotetrameric assembly, which was predicted to form a hollow cage-like assembly comprising 8,864 total amino acids.

**Figure 2.**
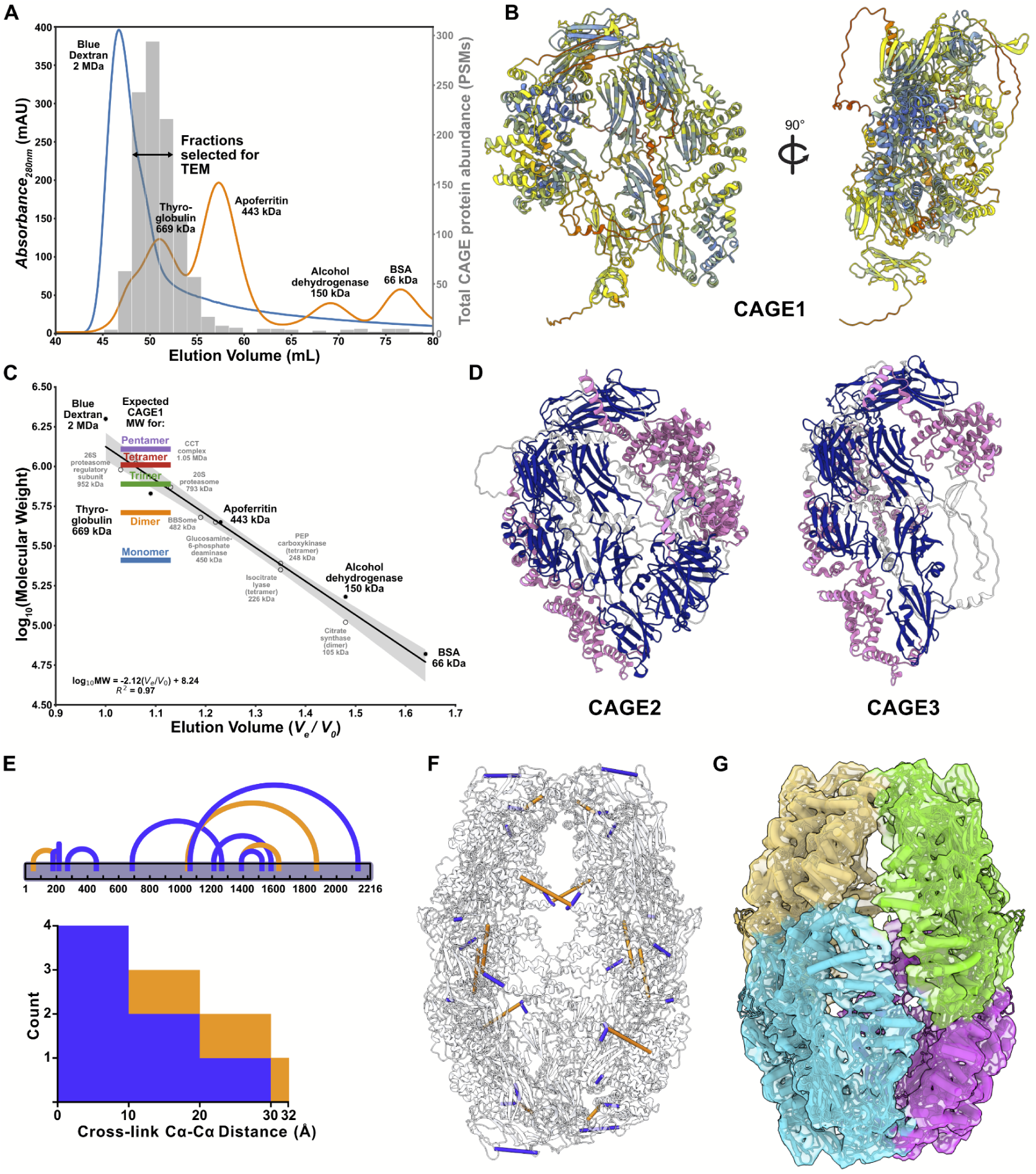
A model of the tetrameric *Tetrahymena* CAGE complex supported by cryo-EM, XL/MS, and structure prediction. **(A)** *Tetrahymena* SEC fractions show the size of the protein complexes imaged by cryo-EM. **(B)** AlphaFold predicted structure of a large abundant “uncharacterized” protein from mass spectrometry data (UniprotID: I7MLD7, CAGE1 protein). **(C)** Calibration of standard protein sizes from SEC to determine possible CAGE protein stoichiometries. **(D)** AlphaFold prediction of *Tetrahymena* CAGE1 paralogs: CAGE2 and CAGE3 (pink, α-helices; navy, β-sheets). **(E)** Intrachain (blue) and interchain (orange) cross-links within the CAGE1 complex all span 32 Å or less, as measured between alpha carbons and plotted onto a tetrameric model of CAGE1 in **(F). (G)** Model of the CAGE1 complex flexibly fit into the cryo-EM density.

### Evidence for homo-tetramers and not mixed heteromers

With I7MLD7_TETTS a compelling candidate for the ellipsoid structure, we searched more deeply across the most abundant proteins in the fraction for other possible alternatives. Notably, two more, large (roughly 2,000 amino acid long) uncharacterized proteins were also evident in the mass spectrometry data, and sequence alignments indicated that these two proteins (I7M9Z0_TETTS and I7MGF4_TETTS) were both homologs of I7MLD7_TETTS. We confirmed by additional sequence searches that these three proteins were likely to be the only members of this sequence family encoded by the *Tetrahymena* genome. Importantly, the peak of the three proteins’ chromatographic elution profiles coincided precisely with the fractions surveyed by electron microscopy, as plotted in **Figure 2A**.

Given these observations and subsequent support described below associating this protein family with the cage-like structure, we assigned new gene symbols to the three *Tetrahymena* paralogs as follows: As ranked from most to least abundant in our proteomics datasets, we opted to call I7MLD7_TETTS (TTHERM_00565630) as CAGE1, I7M9Z0_TETTS (TTHERM_00444610) as CAGE2, and I7MGF4_TETTS (TTHERM_00530680) as CAGE3. The proteins vary by nearly 600 amino acids in length, with CAGE1 of intermediate length (2,216 amino acids long), while CAGE2 is the largest (2,418 amino acids) and CAGE3 the smallest (1,890 amino acids). Nonetheless, AlphaFold predicts all three to adopt the same fold, with the variable ∼600 amino acids being primarily the loss of a portion of the C-terminus in CAGE3, accompanied by small insertions and deletions distributed across the full span of the structures in assorted surface loops and connections between otherwise visually compact domains (**Figure 2D)**.

At this resolution, we were unable to determine whether the observed complexes were exclusively homotetramers or if CAGE paralogs can co-assemble into mixed heterotetramers. However, the structural models and measured protein abundances were both supportive of the dominant form being composed of CAGE1. In support of this, we successfully detected self-cross-links within each CAGE paralog but no cross-links between them, consistent with only homomeric assemblies, as shown for the intramolecular and intermolecular links in CAGE1 (**Figure 2E)**. We also found many CAGE genes existed in single copies in other species (e.g. in the choanoflagellate *Salpingoeca rosetta*), supporting the likelihood of homotetrameric assemblies being a typical form for CAGE complexes.

After identifying our cage-like protein assembly, we fit the AlphaFold predicted structure into the cryo-EM map by rigidly docking 4 copies of the structure into the 3D reconstruction and then flexibly fitting them into the map using NAMDINATOR^26^. In order to validate this model with independent evidence, we mined our mass spectrometry / cross-linking data for intramolecular cross-links that could be mapped onto this model. We found 10 unique amino-acid resolution cross-links, all of which were supported by our model, cross-linking residues separated by 32 Å or less (Cɑ-Cɑ), as expected for the DSSO cross-linker (**Figure 2E-F**). Our consensus model of the *Tetrahymena* CAGE1 complex satisfied both the EM map (**Figure 2G**) and the cross-link distance restraints.

### CAGE exhibits a novel fold and domain organization

We considered the *Tetrahymena* CAGE1 protein as being representative of the overall fold, and next examined its structure and domain organization in some depth. CAGE1 is composed of a set of compact beta-strand domains accompanied by several more extended alpha-helical domains. Searches of the protein domain databases PFAM^27^ and SCOP^28^ failed to identify previously known domains within the protein. Instead, we defined domain boundaries through a combination of visual inspection of the structure and analysis of the boundaries of confidently predicted domains as assessed by AlphaFold’s Predicted Aligned Error (PAE) plots and illustrated in **Figure 3A**. PAE plots have previously been shown to have value for identifying boundaries of folded domains^29–31^, and were particularly useful in the absence of previously identified protein domains.

**Figure 3.**
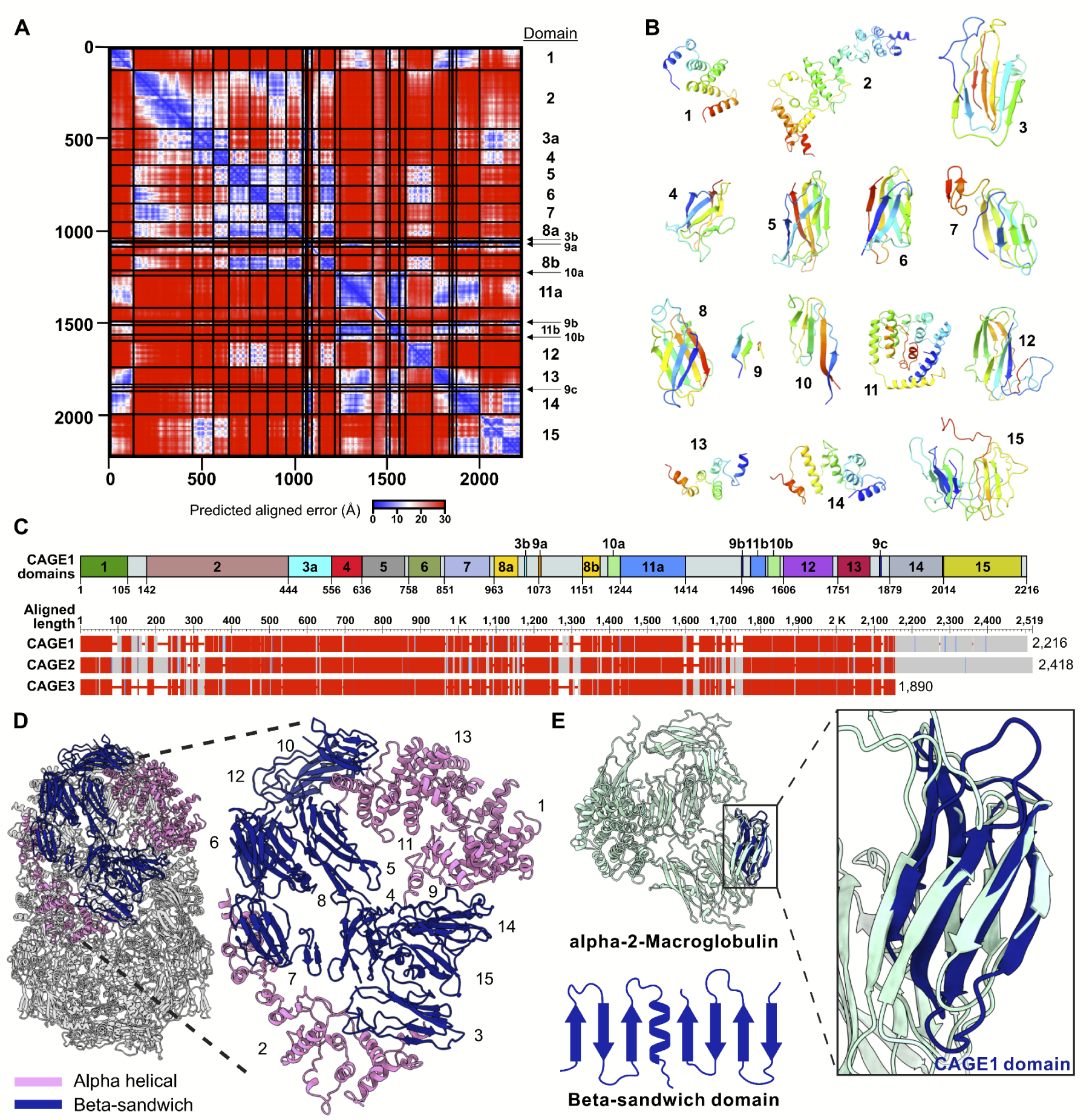
Novel fold and domain architecture of the CAGE complex. **(A)** PAE heatmap of CAGE1 monomer was used to define structural domains within the ∼2,300 aa protein, resulting in 15 domains **(B)**. **(C)** The domains identified in **(A)** mapped onto the CAGE1 primary sequence (top). Alignment of CAGE1 with the two paralogs present in *Tetrahymena*: CAGE2 (UniProt: I7M9Z0) and CAGE3 (UniProt: I7MGF4). **(D)** Assembled CAGE1 tetramer with domains colored pink (α-helical domain) or navy (β-sheet domain). Individual domains from **(B)** are labeled in the assembled structure to highlight the domain packing. **(E)** Alpha-2-macroglobulin subunit shown (light blue) with CAGE domain (navy) identified with Foldseek^54^ with a cartoon diagram of the beta-sandwich domain from the CAGE complex.

In all, we identified 15 distinct domains, defined in order along the linear sequence as follows: domain 1 (amino acids 1-105), domain 2 (142-443), domain 3 (444-555 and 1036-1041), domain 4 (556-635), domain 5 (636-748), domain 6 (758-843), domain 7 (851-951), domain 8 (963-1021,1151-1195), domain 9 (1073-1080, 1496-1500, 1852-1857), domain 10 (1215-1244, 1565-1598), domain 11 (1244-1414, 1519-1559), domain 12 (1606-1738), domain 13 (1751-1833), domain 14 (1879-2011), and domain 15 (2014-2215). As shown in **Figure 3B**, domains 1, 2, 11, 13, and 14 are entirely alpha helical, while the remaining domains form beta sheet domains. The remaining amino acids appear to serve as inter-domain linkers as well as extended, less compact stretches of the protein. Several domains (domains 3, 8, 9, 10, and 11) were composed of non-contiguous amino acid segments, with e.g. multiple strands of the same beta sheet being contributed by amino acid segments distant in the primary chain (**Figure 3C**).

These domains are organized to enclose a hollow central cavity, which presumably represents the site of the (still unknown) CAGE activity. The beta sheet domains colocalize along the structure, while the alpha helical bundles are similarly clustered together (**Figure 3D**). CAGE2 and CAGE3 demonstrate similar organization of beta sheets and alpha helices, with CAGE3 predominantly differing by the absence of domains 14 and 15 (**Figure 2D**).

We used Foldseek^32^ to structurally align each individual domain against the PDB database^33^. While alpha helical domains 1 and 13 (and portions of 11) exhibited similar folds to each other, they showed no matches in the PDB. However, the beta sandwich domains (domains 3, 4, 5, 6, 7, 8) when searched against the PDB, aligned with Immunoglobulin-like domains, either of the Immunoglobulin/Fibronectin type III/E set domains/PapD-like type or the C2 type, as classified by the ECOD database^34^ **(Table S1**), although none of the domains had been previously annotated as such. Interestingly, domains 4 and 6 top hits were alpha-2-macroglobulin, a large glycoprotein that forms a homotetramer with diverse functions. It is perhaps best known for its role as a protease inhibitor where rather than block the active site, the complex contains a bait region and when activated encloses the protease in a cage^35^ (**Figure 3E**).

### The CAGE protein has an unusual phylogenetic distribution across the tree of life

To explore the potential function of the CAGE complex, we searched for homologs using amino acid sequence matching (BLASTP) and 3D structure-structure matching (Foldseek). Unexpectedly, CAGE homologs were not limited to ciliated unicellular eukaryotes but were instead distributed across the full tree of life (**Figure 4A**). Bacterial genes accounted for more than ⅔ of the identified CAGE homologs, and were most prevalent in Pseudomonadati, especially in the PVC superphylum lineages Planctomycetota and Verrucomicrobiota, as well as in other gram negative bacteria such as Thermodesulfobacteria. In contrast, while archaeal homologs were observed, they accounted for only ∼0.1% of the CAGE homologs identified, raising the possibility that the bacterial CAGE homologs might have been horizontally acquired from eukaryotes.

**Figure 4.**
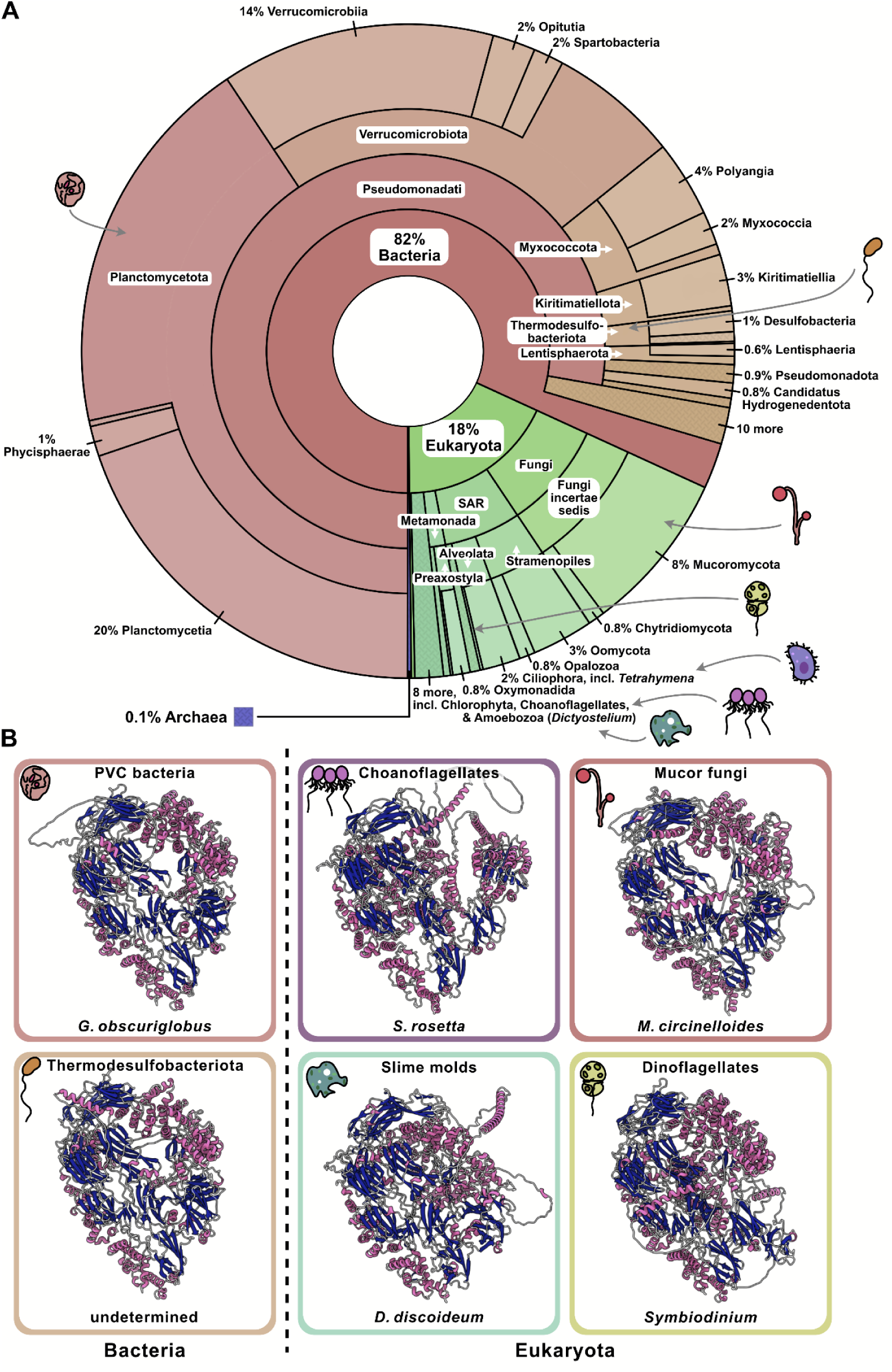
Deep evolutionary conservation of the CAGE complex across gram-negative bacteria and eukaryotic microbes. **(A)** Krona plot^61^ showing the phylogenetic distribution of 3,521 CAGE proteins (ordered radially and labeled with increasing taxonomic specificity towards the perimeter) across the domains of life, with broad representation in gram-negative bacteria and diverse eukaryotes. An interactive version of the plot is available in **File S1**. **(B)** AlphaFold3-predicted structures of representative bacterial and eukaryotic CAGE proteins, colored by secondary structure (pink, α-helices; navy, β-sheets), demonstrate a conserved fold across all species. Most bacterial CAGE genes have been derived from metagenomics sequence assemblies, and, for the case of the Thermodesulfobacteriota CAGE, lack a genus/species assignment.

Within eukaryotes, the CAGE complex was found to be broadly but highly selectively conserved, with homologs found across the ciliates and other SAR organisms, including dinoflagellates (*Symbiodinium*) and stramenopiles (*Oomycota*), but also in plants, such as the green algae *Cymbomonas* and *Volvox* (and present in some *Chlamydomonas* species although absent in *Chlamydomonas reinhardtii*), in Excavata protists (*Strigomonas*, *Angomonas*, and *Phytomonas*), and across Amorphea, spanning choanoflagellates (*Salpingoeca*), fungi (e.g., *Mucor* and *Mortierella*), and amoebozoa (*Dictyostelium*) (**Table S2**). CAGE homologs appeared to be predominantly restricted to unicellular microbes, however, and we were unable to find homologs in animals or flowering plants, despite searching using structure-structure alignments (using Foldseek) or hidden Markov models (using eggNOG/HMMer^36,37^).

To test whether its overall 3D architecture was similarly conserved, we predicted the structures of representative CAGE homologs from both bacteria and eukaryotes using AlphaFold3. Despite the wide phylogenetic spread, the structural domains were strikingly consistent: β-sheets clustered centrally, flanked by α-helices, together forming a conserved concave tertiary arrangement (**Figure 4B**). These results indicate that the CAGE complex has retained its overall fold across at least two billion years of evolution. Given its broad phylogenetic distribution and conserved fold, we next sought to experimentally determine the high-resolution structure of the CAGE complex in a representative non-ciliated eukaryote, *Dictyostelium discoideum*.

### A near-atomic resolution structure of the Dictyostelium CAGE reveals conserved topology

To experimentally confirm that the CAGE complex adopts a conserved architecture across eukaryotic clades, we determined the structure of the CAGE ortholog from the slime mold *Dictyostelium*, as slime molds lack cilia and possess only one CAGE gene (aka AbpF), which exhibits ∼25% sequence identity to the *Tetrahymena* CAGE1 (**Figure 5A**). Using a combination of biochemical fractionation, mass spectrometry, and cryo-EM, we enriched native cell lysates for the endogenous *Dictyostelium* CAGE complex and solved its structure to 3.3 Å resolution (**Figure 5B**).

**Figure 5.**
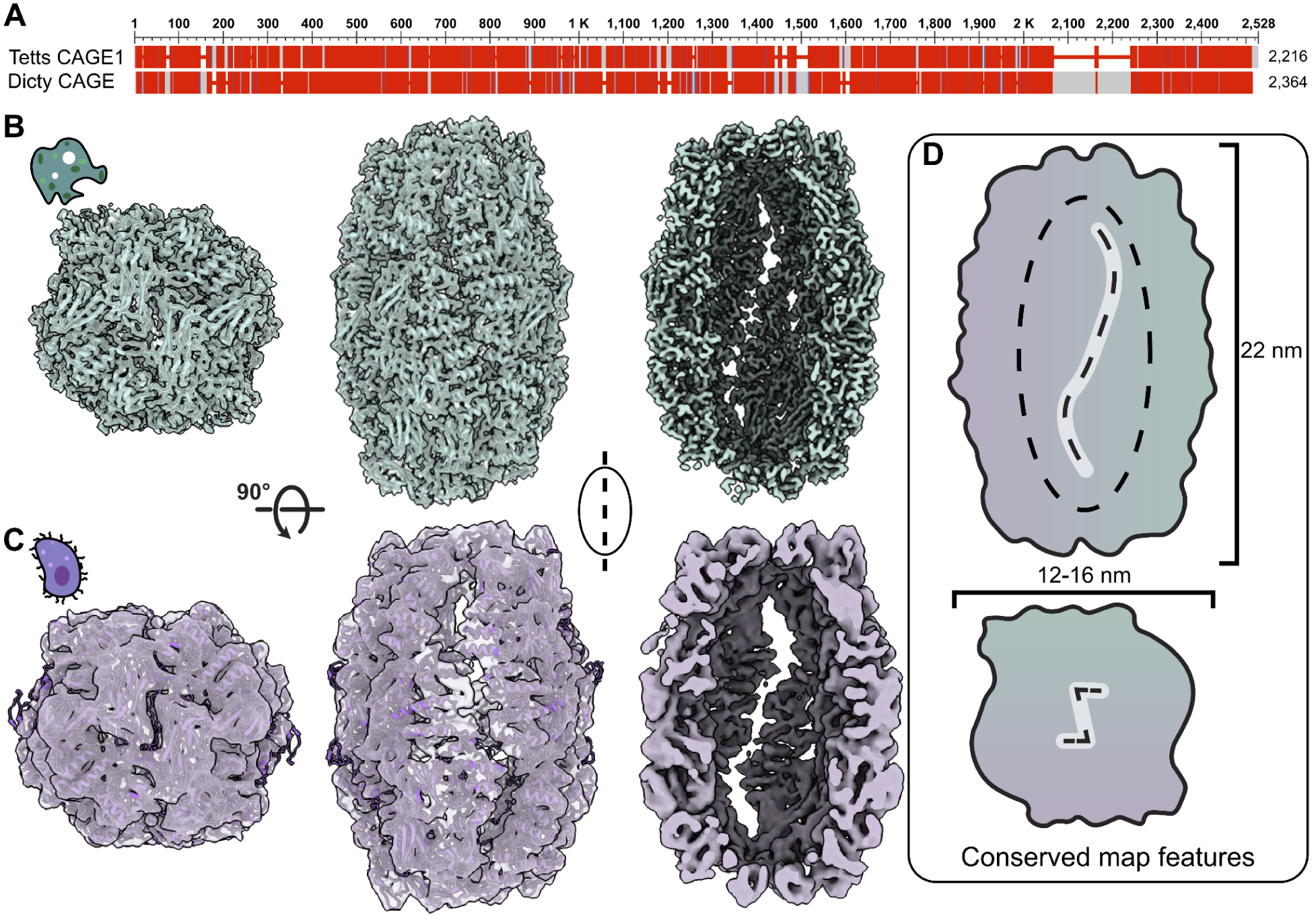
A near-atomic resolution structure of *Dictyostelium* CAGE complex shows major structural features are conserved with *Tetrahymena* CAGE1. **(A)** Pairwise sequence alignment between *Tetrahymena* and *Dictyostelium* CAGE. **(B)** 3.3 Å resolution CAGE structure from cryo-EM of fractionated *Dictyostelium* cellular lysate shows the top view and side view, illustrating the fit of the molecular model into the cryo-EM density. Comparison with the **(C)** structurally-aligned *Tetrahymena* CAGE complex cryo-EM densities and model reveals **(D)** conservation of major structural features: Both CAGE cavities measure about 18 nm in the long axis and have a ∼8 nm diameter enclosing a volume of 600 nm^3^. The side view cutaway reveals a long S-shaped slit and the top view contains a smaller inverted z-shaped opening.

At this resolution, the *Dictyostelium* CAGE displayed striking similarity to the *Tetrahymena* complex (**Figures 5B-D**). Both assemble into a hollow shell ∼22 nm in height and ∼12-16 nm in diameter, enclosing a central cavity that approximates a flattened ellipsoid of 18 x 8 x 8 nm in size, corresponding to approximately 600 nm^3^ in volume. Access to this central cavity is afforded through prominent S-shaped openings on the sides and a backwards Z-shaped opening at the top, architectural features also evident in the *Tetrahymena* structure.

Model building with ModelAngelo^38^ confirmed that the structural domains pack in a highly similar arrangement between *Dictyostelium* and *Tetrahymena*, with the two structures aligning globally with a 10 Å r.m.s.d for the 80% best-fitting residue pairs (1533 Cα pairs), reinforcing the conclusion that CAGE adopts a conserved cage-like topology across diverse eukaryotic clades. This conservation, despite low sequence identity, strongly suggests that the overall fold and cavity architecture are central to the complex’s biological role.

## Discussion

In this study, we identify and structurally characterize the CAGE complex, a previously unknown ∼1 MDa protein assembly conserved across bacteria and eukaryotes. CAGE adopts a novel homotetrameric architecture that encloses a large central cavity, yet lacks obvious conserved enzymatic motifs or cofactors, leaving its biological function unresolved. The deep evolutionary conservation of the CAGE protein family suggests a fundamental biochemical or cell biological role, but one that remains a mystery.

Strikingly, despite our identification of CAGE from *Tetrahymena* cilia, CAGE homologs were found both in ciliated and non-ciliated eukaryotes, e.g. as for ciliated chytrid fungi vs. the non-ciliated mucorales fungi. Even within eukaryotes, the CAGE complex’s broad phylogenetic distribution indicates that it cannot be restricted to only ciliary roles, but likely serves a more general, biochemical or cell biological role. Its extensive conservation across the major eukaryotic clades points to an ancient origin for the complex, likely dating at least to the last common ancestor of eukaryotes and possibly earlier, depending on whether the prokaryotic CAGE genes were vertically or horizontally acquired. Regardless, the many prokaryotic homologs point to a functional role that is conserved in both prokaryotic and eukaryotic contexts, and one that could potentially be readily lost in various eukaryotic lineages, especially multi-cellular ones.

To gain initial clues into possible functional roles, we analyzed intermolecular interactions of the CAGE paralogs in *Tetrahymena* from our XL/MS data. The distribution of these contacts provides some molecular hints toward how CAGE may engage with other cellular proteins. These data revealed numerous interaction partners mapping to defined regions of the CAGE structure, allowing us to predict whether binding occurs on the interior or exterior surfaces of the cage (**Figure 6A**).

**Figure 6.**
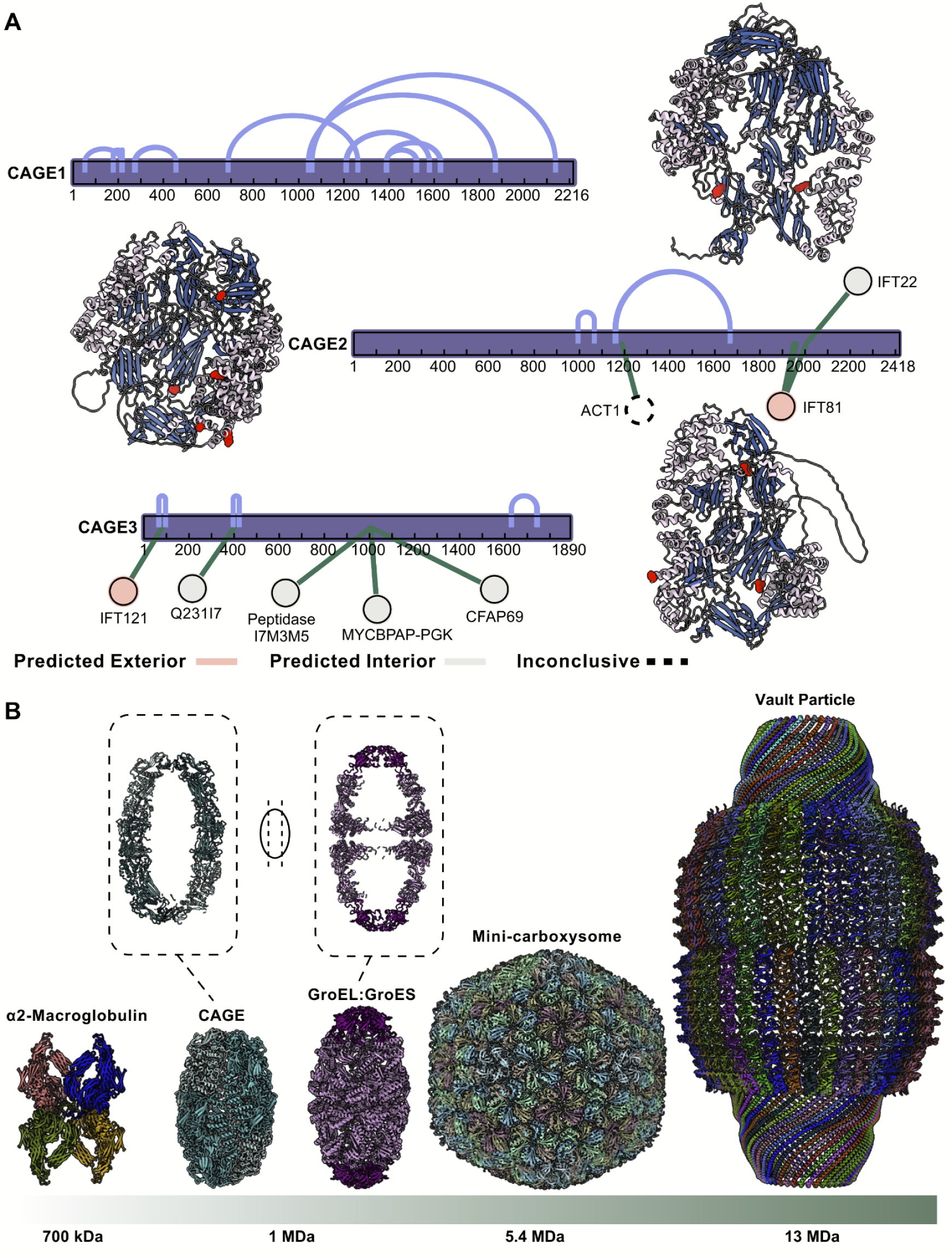
Intermolecular contacts and structural parallels place CAGE among giant molecular assemblies. **(A)** XL/MS identifies interaction partners for the three CAGE paralogs in *Tetrahymena*. Mapping residue-level contacts onto AlphaFold3 predictions reveals their distribution across the protein surface, allowing prediction of whether interactions occur on interior or exterior of the CAGE assembly. **(B)** The CAGE complex (left) shown to scale alongside other large hollow assemblies— human ɑ2 macroglobulin^62^ (PDB: 7O7P, middle), bacterial GroEL:GroES^63^ (PDB: 8BKZ, middle), the bacterial carboxysome^64^ (showing the “mini” carboxysome for easier structural comparison; PDB: 9F0H, middle) and the eukaryotic vault complex^65^ (PDB: 6BP7, right)—illustrating its shared megadalton architecture but distinct morphology.

On the interior of the CAGE we identify contacts with ciliary specific proteins like Cilia And Flagella Associated Protein 69 (CFAP69) and IFT22, in addition to an amino peptidase (I7M3M5), adenylate cyclase (Q231I7), and phosphoglycerate kinase (MYCBPAP-PGK). The exterior of the CAGE makes several contacts with IFT proteins suggesting a possible role in transport. Interestingly, we observe a cross-link to actin, although we cannot determine based on its position whether it is binding the interior or exterior of the CAGE complex. Intriguingly, an interaction with actin was previously reported for the *Dictystelium* CAGE homolog AbpF, from which its name derives (actin-binding protein F)^39^, indicating that this binding activity at least is evolutionarily conserved.

Several additional interaction partners have been reported for other CAGE homologs, including observations of interactions with the histone chaperone Asf1 by the *Tetrahymena* CAGE3 (termed Asf1-interacting protein 1) and CAGE1 (termed Asf1-interacting protein 2) proteins^40^; CAGE2 did not share this activity. However, in our data we do not observe this interaction, and Asf1 is a histone chaperone that apparently lacks homologs in gram negative bacteria. This indicates that CAGE proteins likely interact with multiple partners in a cell type and subcellular context-specific fashion.

A second line of evidence comes from a consideration of subcellular compartments across taxa: using the deep learning model DeepTMHMM^41^, we found that most bacterial CAGE proteins were predicted to have 21-22 amino acid N-terminal secretion signal peptides, within our ability to discriminate full-length CAGE protein sequences derived from metagenome-assembled genomes (MAGs) and scaffolds. This strongly supports prokaryotic CAGEs being transported outside of the cytoplasm, potentially to the periplasmic space or paryphoplasm (a dense, membrane-separated, ribosome-less area at the periphery of *Planctomycetes* cells^42^), as appropriate, or possibly secreted extracellularly or trafficked to endomembrane-bound intracellular compartments, which may be present in *Planctomycetes*, as for example, the anammoxosome organelles that produce gaseous N_2_ from ammonium and nitrite^43^. In contrast, eukaryotic CAGE proteins generally appear to lack detectable secretion signals.

In the absence of more specific functional data, we can at present only speculate about what functions might be performed by CAGE complexes. Although we first identified CAGE inside *Tetrahymena* cilia, the presence of homologs in non-ciliated eukaryotes (slime mold) as well as in prokaryotes, which lack cilia altogether, argues strongly against a ciliary-specific role, especially in light of literature documenting specific CAGE proteins interacting with proteins from other compartments (e.g. actin or histone chaperone binding^39,40^), instead supporting a more general function. One such possibility is a protein or RNA chaperone. It is not uncommon for symmetric, homo-oligomeric assemblies that are broadly phylogenetically distributed across prokaryotes and eukaryotes to act as protein chaperones. This is the case for GroEL/Cpn60 class chaperones, which are, notably, also observed in the same biochemical fractions as the CAGE complex (e.g. **Figure 1**). However, unlike classical chaperonins, we have not yet observed substrate density within the CAGE cavity, and the interior cavity of CAGE is both considerably larger and not compartmentalized in the manner of GroEL/GroES (**Figure 6B**). An alternative is that CAGE represents a new class of molecular container, comparable to alpha-2-macroglobulin^35^, carboxysomes^1^ and vaults^4^, but with a distinct architecture.

Like these other giant assemblies, the discovery of CAGE underscores the power of discovery-driven structural biology to reveal unexpected molecular architectures. The conservation of CAGE across bacteria and eukaryotes suggests that other ancient and enigmatic complexes remain to be uncovered. Pushing *in situ* and survey-based approaches in structural biology to achieve higher resolutions coupled with methodologies like XL/MS may hold the key to important molecular insights. The CAGE complex, thus, stands as both a new structural archetype and a reminder that the molecular universe is far from fully charted.

## Methods

### Tetrahymena culture

*Tetrahymena thermophila* strain SB715 was obtained from the Tetrahymena Stock Center (Cornell University, Ithaca, NY; Washington University in St. Louis) and cultured in Modified Neff Medium (0.25% yeast extract, 0.25% proteose peptone, 0.5% glucose, 33.3 μM FeCl₃) at room temperature (∼22 °C) without shaking. For cilia isolation, cultures were expanded to 3 L and incubated at 30 °C with shaking at 100 rpm.

### Tetrahymena cilia isolation

*Tetrahymena* cells were collected from 3 L of culture by centrifugation (3,000 × g, 5 min, room temperature) and resuspended in 120 mL HNMKS buffer (50 mM HEPES pH 6.9, 36 mM NaCl, 0.1 mM MgSO₄, 1 mM KCl, 250 mM sucrose, 0.1 mM PMSF). Cilia were detached from cells by dibucaine shock (final 1 mg/mL, 3 min), followed by addition of three volumes of HNMKS^44^. Deciliated cell bodies were removed by centrifugation (4,000 × g, 5 min, 4 °C). The isolated cilia pellet was resuspended in HEPES–Cilia Wash Buffer (HEPES-CWB; 50 mM HEPES pH 7.4, 3 mM MgSO₄, 0.1 mM EGTA, 250 mM sucrose, 1 mM DTT, 0.1 mM PMSF, Roche cOmplete Mini EDTA-free protease inhibitors, Roche PhosSTOP EASY phosphatase inhibitors) and lysed with 1% Nonidet P-40 (Roche Applied Science, Cat. #1754599). Axonemes were removed by centrifugation (17,000 × g, 20 min, 4 °C), and the detergent-soluble supernatant is termed the membrane/matrix fraction. Typically yield was ∼2 mL with a protein concentration between 2.3 - 3.1 mg/mL determined by DC Bradford Assay (Bio-Rad).

### Tetrahymena crude CAGE1-containing sample preparation

CAGE1 enrichment from the detergent-soluble membrane/matrix fraction was performed by preparative scale size-exclusion chromatography (SEC). A 2 mL aliquot of soluble extract (∼4.6 - 6.2 mg) in HEPES-CWB was loaded onto a HiLoad 16/600 Superdex 200 PG (preparative grade) column (Cytiva) using a 2 mL sample loop. Chromatography was at a flow rate of 1 mL/min with a mobile phase of 50 mM HEPES pH 7.4, 50 mM NaCl, 3 mM MgSO₄, 0.1 mM EGTA. 1.5 mL fractions were collected and analyzed by mass spectrometry to identify CAGE1-containing fractions. CAGE1 elution positions were benchmarked using a commercial molecular weight standard mixture (Sigma-Aldrich #MWGF1000-1KT; Blue Dextran, bovine thyroglobulin, horse spleen apoferritin, bovine serum albumin, yeast alcohol dehydrogenase) under identical conditions. Fractions 16–18 were pooled (∼ 1 MDa range), clarified by ultracentrifugation (100,000 × g, 1.25 h, 4 °C; Optima MAX-TL) to remove membranous debris (e.g., liposomes observed by EM), and concentrated with a Sartorius Vivaspin Turbo 100,000 MWCO device to 50 μL.

### Dictyostelium strain and culture

*Dictyostelium discoideum* AX2-214 was obtained from the Dicty Stock Center (Columbia University, New York, NY). Frozen cell stocks were thawed and allowed to adhere in a 10 cm dish containing 10 mL HL5 medium (axenic) with 74.8 mM glucose at ∼22 °C for 1 h. Cells were then resuspended in 10 mL HL5 supplemented with 74.8 mM glucose, 10,000 units/mL penicillin, and 10,000 μg/mL streptomycin. Cultures were scaled to 12–16 plates and subsequently expanded to 3 L in shaking flasks (180 rpm, room temperature) to prepare for protein extraction.

### Dictyostelium soluble protein isolation

Cultures were harvested before exceeding a density of 4 × 10⁶ cells/mL. For each 3 L batch, *Dictyostelium* cells were pelleted (500 × g, 4 min, room temperature), washed in 400 mL 50 mM HEPES pH 7.4, pelleted again, and resuspended in 8 mL Dicty Lysis Buffer (50 mM HEPES pH 7.4, 100 mM NaCl, 3 mM MgSO₄, 0.1 mM EGTA, 1 mM DTT, 0.1 mM PMSF, Roche cOmplete Mini EDTA-free protease inhibitors, Roche PhosSTOP EASY phosphatase inhibitors, and 1% Igepal CA-630 (Sigma-Aldrich)). Lysis proceeded on ice for 10 min with brief vortexing each minute. Benzonase was added to 1 U/mL and lysate incubated on ice for 30 min. The lysate was clarified by centrifugation (17,000 × g, 20 min, 4 °C) to remove intact cells, organelles, and insoluble proteins, followed by ultracentrifugation (100,000 × g, 1.25 h, 4 °C) to deplete membranes. The typical yield of detergent-soluble cytosolic proteins was at a concentration of 14–18 mg/mL as measured by DC Bradford (Bio-Rad) prior to SEC.

### Two step Dictyostelium CAGE sample preparation

The *Dictyostelium* sample preparation follows the method described in Hoogerbrugge *et al.* ^45^. CAGE was enriched from the soluble cytosolic samples using preparative scale SEC as for *Tetrahymena* CAGE samples. A 2 mL sample in HEPES–Dicty Lysis Buffer was loaded onto a HiLoad 16/600 Superdex 200 PG (Cytiva) column using a 2 mL sample loop. Chromatography was at a flow rate of 1 mL/min with a mobile phase of 50 mM HEPES pH 7.4, 100 mM NaCl, 3 mM MgSO₄, 0.1 mM EGTA. Fractions (1.5 mL) were collected and analyzed by mass spectrometry to pinpoint CAGE-containing fractions. CAGE elution was estimated using the Sigma-Aldrich #MWGF1000-1KT standard set under identical buffer/flow conditions. High–molecular weight fractions (fractions 16-18) were pooled and concentrated to 50 μL using a Sartorius Vivaspin Turbo 100,000 MWCO concentrator.

Concentrated SEC fractions were buffer-exchanged in an Amicon Ultra 0.5 mL 100,000 MWCO centrifugal filter (Millipore) to reduce ionic strength prior to ion-exchange. Further separation employed a mixed-bed ion-exchange column (PolyLC Inc., #204CTWX0510) on a Dionex UltiMate 3000 HPLC, using a gradient between Buffer A (50 mM HEPES pH 7.4, 3 mM MgSO₄, 0.1 mM EGTA) and Buffer B (50 mM HEPES pH 7.4, 1.5 M NaCl, 3 mM MgSO₄, 0.1 mM EGTA). Fractions of 500 μL were collected into a 96-well plate and analyzed by mass spectrometry. CAGE-containing fractions—corresponding to 500–525 mM NaCl elution—were concentrated and buffer-exchanged to 40 μL using an Amicon Ultra 0.5 mL 100,000 MWCO device (Millipore). Protein yield was typically at a concentration of 1.0-1.3 mg/mL determined by DC Bradford (Bio-Rad).

### Protein mass spectrometry

For the *Tetrahymena* samples, protein identification and quantification were performed on a Thermo Orbitrap Fusion mass spectrometer. Peptides were separated by reverse phase chromatography on a Dionex Ultimate 3000 RSLCnano UHPLC system (Thermo Scientific) configured with a C18 trap to Acclaim C18 PepMap RSLC column (Dionex; Thermo Scientific). Elution was with a 3%–45% gradient over 60 min. Peptides were directly injected via a nano-electrospray for data-dependent tandem mass spectrometry ^46^. The mass spectra were processed with the Proteome Discoverer (Thermo Fisher) standard workflow as described below.

Raw mass spectra files were analyzed in Proteome Discoverer 2.3 against the *Tetrahymena* proteome (26,976 entries; UniProt, 2018) plus a 379-protein contaminant database ^47^. Searches settings were for trypsin specificity with up to two missed cleavages, a precursor mass range of 350–5,000 Da, and a mass tolerance of 10 ppm, dynamic N-terminal modifications of acetylation (+42.011 Da), methionine loss (−131.040 Da), or methionine loss followed by acetylation (−89.030 Da), static modification of carbamidomethyl (+57.021 Da), and protein grouping by maximum parsimony. Peptide-level false discovery rate (FDR) was controlled at <1% (strict); protein-level FDR thresholds were <1% (strict) and <5% (relaxed).

For *Dictyostelium* samples, protein identification and quantification were performed on a Thermo Orbitrap Fusion Lumos Tribrid mass spectrometer. Peptides were separated by reverse phase chromatography on a Dionex UltiMate 3000 RSLCnano UHPLC (Thermo Scientific) configured with a C18 trap coupled to an EASY Spray PepMap RSLC C18 analytical column (Thermo Scientific). The mass spectra were acquired using a standard top-speed HCD MS1-MS2 method with a 60 minute gradient and stepped HCD and processed with the Proteome Discoverer (Thermo Fisher) standard workflow as described below.

Raw mass spectra were processed in Proteome Discoverer 2.3 against the UniProt *Dictyostelium* proteome (12,726 entries; January 2024) supplemented with a 379-protein contaminant set from the Hao group ^47^. Searches used the same parameters as above.

### Cryogenic electron microscopy

#### Tetrahymena

SEC-enriched, pooled fractions were adjusted to have a final concentration of 50 mM HEPES pH 7.4, 100 mM NaCl, 3 mM MgSO_4_, 0.1mM EGTA, 1 mM DTT, and 5% glycerol to reduce salt concentration and include a cryoprotectant for plunge-freezing. A 3 µL aliquot was applied to a Quantifoil Graphene Oxide (GO) 2/4 Cu200 mesh, maintained at 100% humidity and 4 °C. We waited for 60 seconds before blotting the sample of a blot force of 1 for 3 seconds, then plunged it into liquid ethan using an FEI Vitrobot Mark IV (Thermo Fisher Scientific). A total of 22,764 micrographs were collected using a 200 kV FEI Glacios (Thermo Fisher Scientific) cryo-transmission electron microscope operating at 200 kV, equipped with a Falcon IV direct electron detector (Thermo Fisher Scientific). Exposures were collected with a calibrated pixel size of 0.94 Å/pixel, a total electron dose of 50 e−/Å² with a total exposure time of 10 s, and a defocus range between -1.5 and -2.5 µm.

#### Dictyostelium

Concentrated fractions from mixed-bed ion-exchange chromatography were adjusted to a final composition of 50 mM HEPES (pH 7.4), 100 mM NaCl, 3 mM MgSO₄, 0.1 mM EGTA, 1 mM DTT, 0.1 mM PMSF, Roche cOmplete Mini EDTA-free protease inhibitors, Roche PhosSTOP EASY phosphatase inhibitors, and 2% glycerol. This adjustment was made to lower the salt concentration and incorporate a cryoprotectant for subsequent plunge-freezing. A 3 µL aliquot of the concentrated protein solution was applied to C-Flat Holey Thick 40 nm Carbon 1.2/1.3 Cu400 mesh grids coated with GO. For GO coating, C-Flat grids were glow-discharged (60 s), coated with 0.2 mg/mL GO on the carbon side for 1 min, wicked with filter paper, washed three times with deionized water, and air-dried 12–16 h at room temperature. Sample vitrification was performed using an FEI Vitrobot Mark IV (Thermo Fisher Scientific). After sample application and a wait time of 60s at 100% humidity 4 °C, the sample was blotted for 3 s with a blot force of 1 and plunged into liquid ethane. A total of 5,183 micrographs were collected on a 200 kV FEI Glacios cryo-transmission electron microscope operating at 200 kV (Thermo Fisher Scientific) equipped with a Falcon IV direct electron detector (Thermo Fisher Scientific). Exposures were collected with a calibrated pixel size of 0.933 Å/pixel, a total electron dose of 49 e−/Å², and a defocus range between -1.5 and -2.5 µm.

GO support was essential for capturing CAGE particles from both *Tetrahymena* and *Dictyostelium*; no CAGE particles were evident in earlier analyses using uncoated C-Flat Holey Thick 40 nm Carbon 1.2/1.3 Cu400 grids^45^.

### Cryo-EM data processing, building, and refinement

#### General

Map resolution was evaluated using the cryoSPARC Validation (FSC) tool applying the gold-standard Fourier Shell Correlation (FSC) criterion with a 0.143 threshold, and additional map quality metrics were obtained with Phenix cryo-EM validation tools^48^. Processing schemes are shown in **Figure S1** (*Tetrahymena*) and **Figure S2** (*Dictyostelium*), with particle counts, FSC thresholds, and final resolutions summarized in **Table S3**.

#### Tetrahymena

Each dataset was processed in cryoSPARC v4/v4.7^22^, including motion correction, defocus estimation, and micrograph curation. Of the 22,764 acquired micrographs, 17,886 were retained for further analysis, with exclusions largely due to excessive ice thickness or beam-induced drift. An initial set of 44 particles was manually picked, and 2D classification of these manually picked particles was used to generate a template for subsequent template-based particle picking. Particles selected from 2D classes were first subjected to *ab initio* 3D reconstruction, and each *ab initio* class was further refined by heterogeneous refinement with no symmetry applied (C1) for both steps. Particle alignments from the best heterogeneous-refinement class were then refined by non-uniform (NU) refinement in C1, followed by an additional round of NU refinement with D2 symmetry applied. To further refine the particle stack, an additional D2-symmetry heterogeneous refinement with two output classes was performed, and the best class was carried forward to a final NU refinement in D2 symmetry. The final map reached a nominal resolution of 5.64 Å from 3,656 particles (**Figure S1**).

#### Dictyostelium

On-the-fly processing was conducted in cryoSPARC Live ^22^ encompassing motion correction, defocus estimation, micrograph curation, and particle picking. Downstream particle extraction, stack curation, and refinements were performed in cryoSPARC v4/v4.5^22^. Of 5,183 micrographs, 1,319 were retained for further analysis; removals were primarily attributable to thick ice or drift. We used the *Tetrahymena* CAGE1 map as a reference for template-based particle picking. Particles were first processed through *ab initio* 3D reconstruction and homogeneous refinement in C1. Afterward, particles were re-extracted with a reduced box size and re-aligned using homogeneous refinement in C1. The same refinement job was then run with an applied D2 symmetry. Homogeneous refinement with D2 symmetry was then introduced. Following symmetry application, particles were re-extracted using a larger box, duplicate picks were eliminated, and a final homogeneous refinement was run with D2 symmetry imposed. The final map, reconstructed to a nominal resolution of 3.30 Å, was generated from a curated subset of 5,565 particles.

### Model building and refinement

#### Tetrahymena

The CAGE1 structure was predicted using AlphaFold2^24^ and then rigidly docked into the cryo-EM map. After rigid docking, NAMDINATOR^26^ was used to flexibly fit the remainder of the structure into the cryo-EM map, with a final refinement using PHENIX real-space refine^49^. Intramolecular cross-links were mapped onto the flexibly fit structure as additional validation.

#### Dictyostelium

The final refined map was used for input into ModelAngelo^38^ with the CAGE protein sequence. The resulting structure was then further refined within the monomeric boundary of the asymmetric unit of the complex. Four copies of the final monomeric structure were then fitted into the cryo-EM map and refined using PHENIX real-space refine^49^. Coiled regions of the protein chain connecting compact domains were still somewhat unresolved in both the *Tetrahymena* and *Dictyostelium* models, so we could not reject the possibility of 3D domain swapping between the monomeric subunits in either^50^. The deposited *Dictyostelium* chain assignments reflect a reassignment of residues 163-1741 between the A/B chains and similarly between the C/D chains, in order to assign ModelAngelo-built domains (separated by unmodeled gaps) to chains in a manner compatible with the observed tetrameric packing, supporting 3D domain swapping in the tetramer relative to the AlphaFold-predicted monomer structures.

### Cross-linking mass spectrometry

Cross-linking mass spectrometry data were previously collected for independent separations of the corresponding *Tetrahymena* biochemical fractions and are described in full in McCafferty *et al*.^23^. Briefly, aliquots of SEC fractions (approx. 40 ug of protein) that were imaged by cryo-EM were cross-linked by the addition of 5 mM DSSO. Tandem mass spectra (MS/MS^2^/MS^3^) were collected on a Thermo Orbitrap Fusion Lumos tribrid mass spectrometer, as previously described. Data was analyzed using the XlinkX node of Proteome Discoverer 2.3^51^. The full set of cross-links can be found in the following resource article^52^.

### Sequence and phylogenetic analyses

Homologs of the *Tetrahymena* and *Dictyostelium* CAGE proteins were identified by sequence-sequence matching using NCBI BLASTP versus the May 21, 2025 NCBI nr_clustered database^53^, requiring E-value ≤ 10^-5^ and >5% query sequence coverage. **Table S2** provides the full set of 3,521 BLASTP hits obtained. We conducted a parallel search using structure-structure matching with Foldseek to search available databases of predicted AlphaFold and ESMfold structures^54^ and confirmed that hits by the two methods were consistent. The Krona plot of **Figure 4** was generated in turn from this set of homologs using the Krona Excel template method (available from https://github.com/marbl/Krona/wiki/ExcelTemplate). **Figure S3** provides an unrooted maximum likelihood phylogenetic tree of the 1,000 top-scoring CAGE homologs, computed from the BLAST pairwise alignments using Fast Minimum Evolution^55^, a maximal allowable fraction of mismatches of 0.85, and the Grishin (protein) distance criterion^56^, and visualized using iTOL^57^.

## Acknowledgements

The authors gratefully acknowledge the generous support of the Tetrahymena stock center (Cornell University, Ithaca, NY, and Washington University in St. Louis, MO) and the Dicty Stock Center (DSC) at Columbia University, NY. Research was funded by grants from the National Institute of General Medical Sciences R35GM122480 (to E.M.M.) and R35GM138348 (to D.W.T.), National Science Foundation (2019238253 to C.L.M.), National Institute of Child Health and Human Development (HD085901 to E.M.M.), Army Research Office (W911NF-12-1-0390 to E.M.M.), and Welch Foundation (F-1515 to E.M.M., F-1938 to D.W.T.). D.W.T. is a CPRIT Scholar supported by Cancer Prevention and Research Institute of Texas (RR160088). The authors acknowledge the Texas Advanced Computing Center at The University of Texas at Austin for providing high-performance computing resources that contributed to the research results reported in this paper. Cryo-TEM was performed at the University of Texas at Austin Sauer Structural Biology Laboratory (RRID:SCR_022951).

## Authors’ contributions

C.L.M.: Conceptualization, Methodology, Formal analysis, Investigation, Visualization, Resources, Supervision, Writing—original draft, Writing—review and editing.

G.H.: Methodology, Validation, Formal analysis, Investigation, Visualization, Writing—original draft, Writing—review and editing.

O.P.: Methodology, Formal analysis, Investigation.

E.A.S: Formal analysis.

S.R.: Investigation.

D.W.T: Resources; Supervision.

A.F.B.: Formal analysis, Investigation, Resources.

E.M.M.: Conceptualization, Methodology, Formal analysis, Investigation, Visualization, Resources, Supervision, Writing—original draft, Writing—review and editing.

## Data availability

Mass spectrometry proteomics data were deposited in the MassIVE/ProteomeXchange database^58^ under accession numbers MSV000099251/ PXD068692, respectively (*Tetrahymena*) and MSV000099249/ PXD068690, respectively (*Dictyostelium*). Cryo-electron microscopy data was deposited in the Electron Microscopy Data Bank^59^ under accession numbers EMD-75779 (*Tetrahymena*) and EMD-75780 (*Dictyostelium*). Coordinates for the Tetrahymena CAGE1 complex and the Dictyostelium CAGE complex were deposited in the Protein Data Bank^60^ under accession numbers 11KS (*Tetrahymena*) and 11KT (*Dictyostelium*). Prior to peer review, all data is available in a Zenodo repository at doi: 10.5281/zenodo.18925749, including model coordinates, **Figures S1-3**, **Tables S1-3**, and an interactive version of the Krona plot in **Figure 4A (File S1)**.

## Additional Supporting Data and Tables

### Supplemental Figures

**Figure S1** - Cryo-EM workflow and statistics for *Tetrahymena* CAGE1 complex

**Figure S2** - Cryo-EM workflow and statistics for *Dictyostelium* CAGE complex

**Figure S3** - Unrooted maximum likelihood phylogenetic tree of CAGE protein homologs

### Supplemental Tables

**Table S1** - ECOD domain identifications

**Table S2** - CAGE homologs with BLASTP scores and taxonomy

**Table S3** - Cryo-EM statistics

### Supplemental File

**File S1** - Interactive Krona plot.html

## References

1. Shively, J. M., Ball, F., Brown, D. H. & Saunders, R. E. Functional Organelles in Prokaryotes: Polyhedral Inclusions (Carboxysomes) of Thiobacillus neapolitanus. Science 182, 584–586 (1973).

2. Liu, J.-L. Intracellular compartmentation of CTP synthase in Drosophila. J Genet Genomics 37, 281–296 (2010).

3. Kedersha, N. L. & Rome, L. H. Preparative agarose gel electrophoresis for the purification of small organelles and particles. Anal Biochem 156, 161–170 (1986).

4. Kedersha, N. L. & Rome, L. H. Isolation and characterization of a novel ribonucleoprotein particle: large structures contain a single species of small RNA. J Cell Biol 103, 699–709 (1986).

5. McCafferty, C. L., Verbeke, E. J., Marcotte, E. M. & Taylor, D. W. Structural Biology in the Multi-Omics Era. J Chem Inf Model 60, 2424–2429 (2020).

6. Maco, B. et al. Proteomic and electron microscopy survey of large assemblies in macrophage cytoplasm. Mol Cell Proteomics 10, M111 008763 (2011).

7. Kastritis, P. L. et al. Capturing protein communities by structural proteomics in a thermophilic eukaryote. Mol Syst Biol 13, 936 (2017).

8. Ho, C.-M. et al. Bottom-up structural proteomics: cryoEM of protein complexes enriched from the cellular milieu. Nat Methods 17, 79–85 (2020).

9. Su, C.-C. et al. A ‘Build and Retrieve’ methodology to simultaneously solve cryo-EM structures of membrane proteins. Nat Methods 18, 69–75 (2021).

10. Kim, G., Jang, S., Lee, E. & Song, J.-J. EMPAS: Electron Microscopy Screening for Endogenous Protein Architectures. Mol Cells 43, 804–812 (2020).

11. Kirykowicz, A. M. & Woodward, J. D. Shotgun EM of mycobacterial protein complexes during stationary phase stress. Current Research in Structural Biology 2, 204–212 (2020).

12. Kyrilis, F. L., Meister, A. & Kastritis, P. L. Integrative biology of native cell extracts: a new era for structural characterization of life processes. Biological Chemistry 400, 831–846 (2019).

13. Mitchell, D. R. Evolution of Cilia. Cold Spring Harb Perspect Biol 9, a028290 (2017).

14. Sae-Lee, W. et al. The protein organization of a red blood cell. Cell Reports 40, 111103 (2022).

15. Boldt, K. et al. An organelle-specific protein landscape identifies novel diseases and molecular mechanisms. Nat Commun 7, 11491 (2016).

16. Yuan, S. & Sun, Z. Expanding Horizons: Ciliary Proteins Reach Beyond Cilia. Annu. Rev. Genet. 47, 353–376 (2013).

17. Piperno, G., Huang, B. & Luck, D. J. Two-dimensional analysis of flagellar proteins from wild-type and paralyzed mutants of Chlamydomonas reinhardtii. Proc. Natl. Acad. Sci. U.S.A. 74, 1600–1604 (1977).

18. Blackburn, K., Bustamante-Marin, X., Yin, W., Goshe, M. B. & Ostrowski, L. E. Quantitative Proteomic Analysis of Human Airway Cilia Identifies Previously Uncharacterized Proteins of High Abundance. J. Proteome Res. 16, 1579–1592 (2017).

19. Pigino, G. et al. Comparative structural analysis of eukaryotic flagella and cilia from Chlamydomonas, Tetrahymena, and sea urchins. Journal of Structural Biology 178, 199–206 (2012).

20. Bertiaux, E. et al. The Luminal Ring Protein C2CD3 Acts as a Radial In-to-Out Organizer of the Distal Centriole and Appendages. Preprint at 10.1101/2025.06.17.660204 (2025).

21. Chen, Z. et al. In situ cryo-electron tomography reveals the asymmetric architecture of mammalian sperm axonemes. Nat Struct Mol Biol 30, 360–369 (2023).

22. Punjani, A., Rubinstein, J. L., Fleet, D. J. & Brubaker, M. A. cryoSPARC: algorithms for rapid unsupervised cryo-EM structure determination. Nat Methods 14, 290–296 (2017).

23. McCafferty, C. L. et al. Integrative modeling reveals the molecular architecture of the intraflagellar transport A (IFT-A) complex. Elife 11, e81977 (2022).

24. Jumper, J. et al. Highly accurate protein structure prediction with AlphaFold. Nature 596, 583–589 (2021).

25. Evans, R., et al. Protein complex prediction with AlphaFold-Multimer. bioRxiv DOI:10.1101/2021.10.04.463034 2021.10.04.463034 (2022) doi:10.1101/2021.10.04.463034.

26. Kidmose, R. T. et al. Namdinator - automatic molecular dynamics flexible fitting of structural models into cryo-EM and crystallography experimental maps. IUCrJ 6, 526–531 (2019).

27. Mistry, J. et al. Pfam: The protein families database in 2021. Nucleic Acids Res 49, D412–D419 (2021).

28. Andreeva, A., Kulesha, E., Gough, J. & Murzin, A. G. The SCOP database in 2020: expanded classification of representative family and superfamily domains of known protein structures. Nucleic Acids Res 48, D376–D382 (2020).

29. McCafferty, C. L., Pennington, E. L., Papoulas, O., Taylor, D. W. & Marcotte, E. M. Does AlphaFold2 model proteins’ intracellular conformations? An experimental test using cross-linking mass spectrometry of endogenous ciliary proteins. Commun Biol 6, 421 (2023).

30. Zhang, J., Schaeffer, R. D., Durham, J., Cong, Q. & Grishin, N. V. DPAM: A domain parser for AlphaFold models. Protein Science 32, e4548 (2023).

31. Lau, A. M., Kandathil, S. M. & Jones, D. T. Merizo: a rapid and accurate protein domain segmentation method using invariant point attention. Nat Commun 14, 8445 (2023).

32. Van Kempen, M. et al. Fast and accurate protein structure search with Foldseek. Nat Biotechnol 10.1038/s41587-023-01773-0 (2023) doi:10.1038/s41587-023-01773-0.

33. van Kempen, M. et al. Fast and accurate protein structure search with Foldseek. Nat Biotechnol 42, 243–246 (2024).

34. Cheng, H. et al. ECOD: An Evolutionary Classification of Protein Domains. PLOS Computational Biology 10, e1003926 (2014).

35. Rehman, A. A., Ahsan, H. & Khan, F. H. Alpha-2-macroglobulin: A physiological guardian. Journal Cellular Physiology 228, 1665–1675 (2013).

36. Potter, S. C. et al. HMMER web server: 2018 update. Nucleic Acids Research 46, W200–W204 (2018).

37. Huerta-Cepas, J. et al. eggNOG 5.0: a hierarchical, functionally and phylogenetically annotated orthology resource based on 5090 organisms and 2502 viruses. Nucleic Acids Research 47, D309–D314 (2019).

38. Jamali, K. et al. Automated model building and protein identification in cryo-EM maps. Nature 628, 450–457 (2024).

39. Niebling, K. R. Purification and characterization of a 270 kilodalton actin-binding protein from Dictyostelium that localizes to the centrosome. (1993).

40. Garg, J. et al. Conserved Asf1–importin β physical interaction in growth and sexual development in the ciliate Tetrahymena thermophila. Journal of Proteomics 94, 311–326 (2013).

41. Hallgren, J. et al. DeepTMHMM predicts alpha and beta transmembrane proteins using deep neural networks. Preprint at 10.1101/2022.04.08.487609 (2022).

42. Lindsay, M. et al. Cell compartmentalisation in planctomycetes: novel types of structural organisation for the bacterial cell. Archives of Microbiology 175, 413–429 (2001).

43. Fuerst, J. A. & Sagulenko, E. Beyond the bacterium: planctomycetes challenge our concepts of microbial structure and function. Nat Rev Microbiol 9, 403–413 (2011).

44. Dentler, W. L. Chapter 55 Isolation and Fractionation of Ciliary Membranes from Tetrahymena. in Methods in Cell Biology vol. 47 397–400 (Elsevier, 1995).

45. Hoogerbrugge, G., Keatinge-Clay, A. T. & Marcotte, E. M. Serendipity and the slime mold: a visual survey of high molecular-weight protein assemblies reveals the structure of the polyketide synthase Pks16. Mol Cell Proteomics 101484 (2025) doi:10.1016/j.mcpro.2025.101484.

46. McWhite, C. D. et al. A Pan-plant Protein Complex Map Reveals Deep Conservation and Novel Assemblies. Cell 181, 460–474.e14 (2020).

47. Frankenfield, A. M., Ni, J., Ahmed, M. & Hao, L. Protein Contaminants Matter: Building Universal Protein Contaminant Libraries for DDA and DIA Proteomics. J. Proteome Res. 21, 2104–2113 (2022).

48. Liebschner, D. et al. Macromolecular structure determination using X-rays, neutrons and electrons: recent developments in *Phenix*. Acta Crystallogr D Struct Biol 75, 861–877 (2019).

49. Afonine, P. V. et al. Real-space refinement in *PHENIX* for cryo-EM and crystallography. Acta Crystallogr D Struct Biol 74, 531–544 (2018).

50. Bennett, M. J., Schlunegger, M. P. & Eisenberg, D. 3D domain swapping: A mechanism for oligomer assembly. Protein Science 4, 2455–2468 (1995).

51. Liu, F., Lössl, P., Scheltema, R., Viner, R. & Heck, A. J. R. Optimized fragmentation schemes and data analysis strategies for proteome-wide cross-link identification. Nat Commun 8, 15473 (2017).

52. McCafferty, C. L. et al. An amino acid-resolution interactome for motile cilia identifies the structure and function of ciliopathy protein complexes. Developmental Cell 60, 965–978.e3 (2025).

53. Sayers, E. W. et al. Database resources of the National Center for Biotechnology Information. Nucleic Acids Res 52, D33–D43 (2024).

54. van Kempen, M. et al. Fast and accurate protein structure search with Foldseek. Nat Biotechnol 10.1038/s41587-023-01773-0 (2023) doi:10.1038/s41587-023-01773-0.

55. Desper, R. & Gascuel, O. Theoretical foundation of the balanced minimum evolution method of phylogenetic inference and its relationship to weighted least-squares tree fitting. Mol Biol Evol 21, 587–598 (2004).

56. Grishin, N. V. Estimation of the number of amino acid substitutions per site when the substitution rate varies among sites. J Mol Evol 41, 675–679 (1995).

57. Letunic, I. & Bork, P. Interactive Tree Of Life (iTOL) v5: an online tool for phylogenetic tree display and annotation. Nucleic Acids Res 49, W293–W296 (2021).

58. Deutsch, E. W. et al. The ProteomeXchange consortium in 2020: enabling ‘big data’ approaches in proteomics. Nucleic Acids Research gkz984 (2019) doi:10.1093/nar/gkz984.

59. The wwPDB Consortium et al. EMDB—the Electron Microscopy Data Bank. Nucleic Acids Research 52, D456–D465 (2024).

60. Burley, S. K. et al. RCSB Protein Data Bank (RCSB.org): delivery of experimentally-determined PDB structures alongside one million computed structure models of proteins from artificial intelligence/machine learning. Nucleic Acids Research 51, D488–D508 (2023).

61. Ondov, B. D., Bergman, N. H. & Phillippy, A. M. Interactive metagenomic visualization in a Web browser. BMC Bioinformatics 12, 385 (2011).

62. Luque, D. et al. Cryo-EM structures show the mechanistic basis of pan-peptidase inhibition by human α_2_ -macroglobulin. Proc. Natl. Acad. Sci. U.S.A. 119, e2200102119 (2022).

63. Torino, S., Dhurandhar, M., Stroobants, A., Claessens, R. & Efremov, R. G. Time-resolved cryo-EM using a combination of droplet microfluidics with on-demand jetting. Nat Methods 20, 1400–1408 (2023).

64. Wang, P. et al. Molecular principles of the assembly and construction of a carboxysome shell. Sci. Adv. 10, eadr4227 (2024).

65. Ding, K. et al. Solution Structures of Engineered Vault Particles. Structure 26, 619–626.e3 (2018).

