## Supplemental Figures for "The CAGE complex: a hollow, megadalton, protein assembly in prokaryotic and eukaryotic microbes"

### Supplemental Figure Legends

**Figures S1-S2. Cryo-EM workflow and statistics for (S1) *Tetrahymena* CAGE1 complex and (S2) *Dictyostelium* CAGE complex.** In each case, the specific cryo-EM workflow is presented, along with a view of the final EM density and the statistics of the final reconstruction. (A) Data processing and refinement pipeline for the complex using cryoSPARC v4/v4.7 (*Tetrahymena*) and v4/4.5 (*Dictyostelium*). The symmetry applied at each refinement step is indicated at the lower right of the respective refinement. (B) Cryo-EM volume colored by local resolution estimation (units in Å). (C) Gold-standard Fourier shell correlation (GSFSC) curves calculated in cryoSPARC, with the 0.143 threshold. (D) Viewing angle distribution plot.

**Figure S3. Unrooted maximum likelihood phylogenetic tree of CAGE protein homologs.** Calculated based on NCBI BLASTP of the *Dictyostelium* AbpF amino acid sequence versus the nr_clustered database, selecting the top-scoring 1,000 hits, all of which exhibited an E-value ≤ $10^{-50}$.

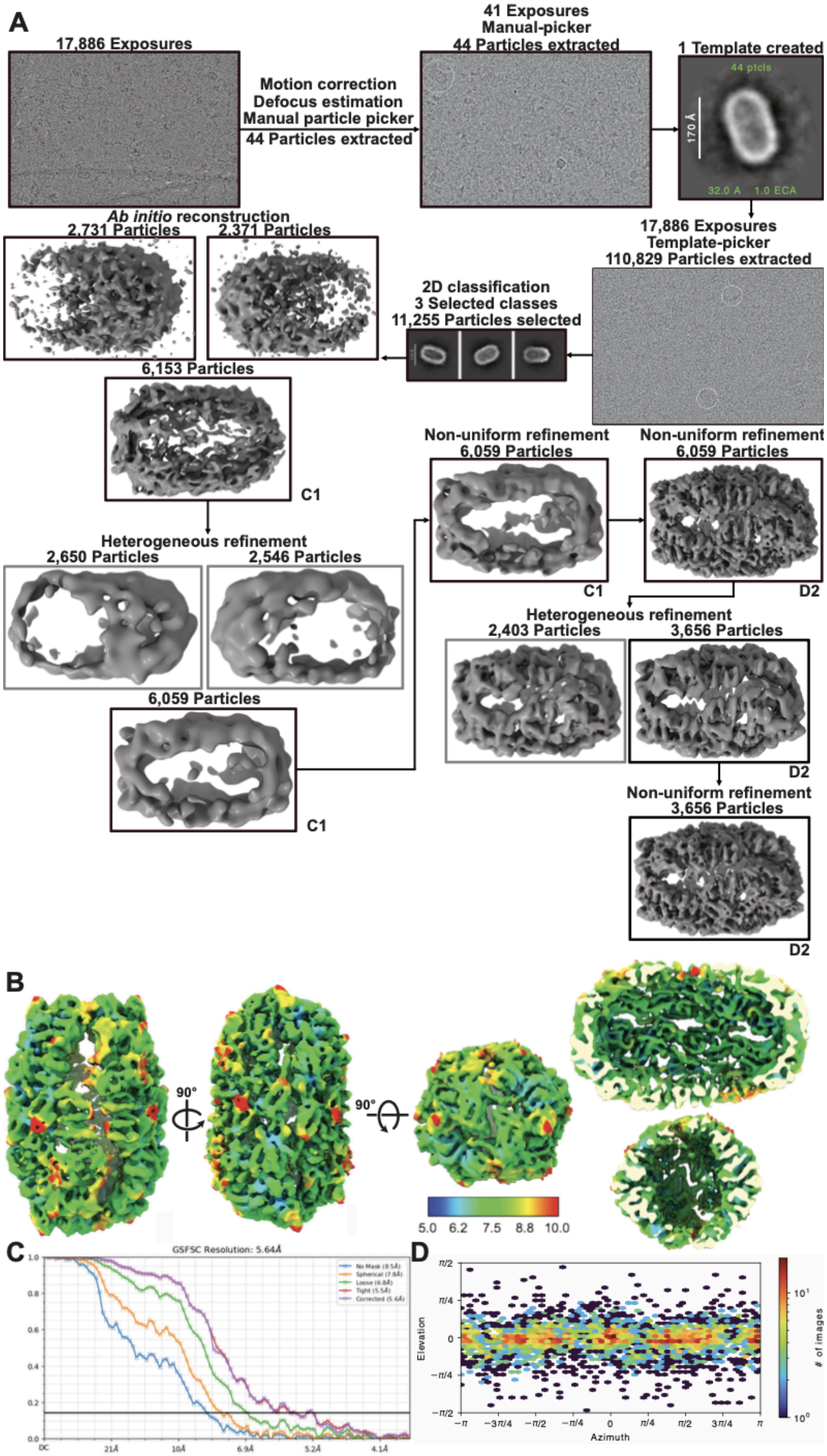


**Figure S1. Cryo-EM workflow and statistics for *Tetrahymena* CAGE1 complex.**

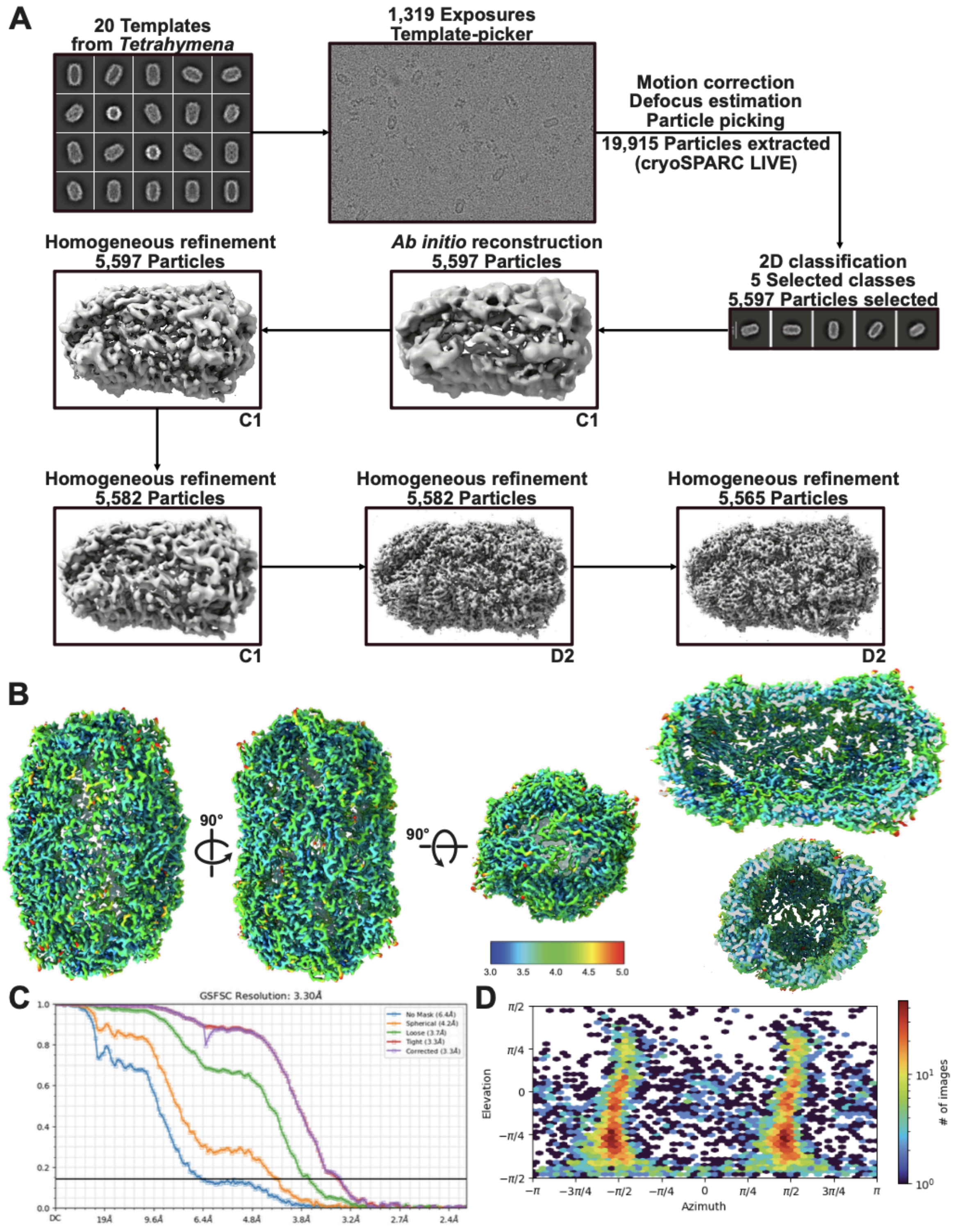


**Figure S2. Cryo-EM workflow and statistics for *Dictyostelium* CAGE complex.**

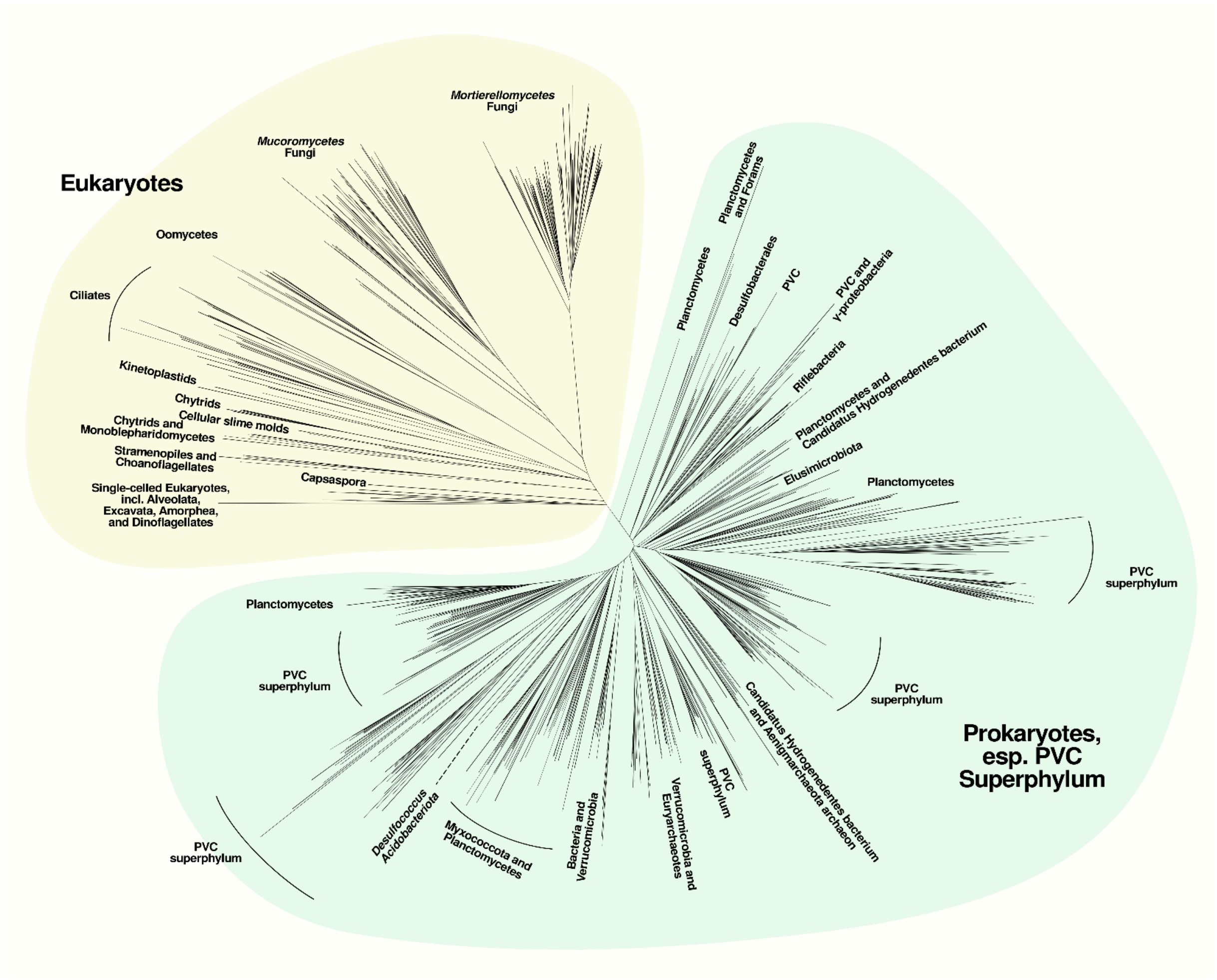


**Figure S3. Unrooted maximum likelihood phylogenetic tree of CAGE protein homologs.**
