## Supplementary material for "The CAGE complex: a hollow, megadalton, protein assembly in prokaryotic and eukaryotic microbes": File S1: File S1 - Interactive Krona plot.html

Javascript must be enabled to view this page.

magnitude
score


CAGE complex

 3521

 1

 638

 1

 14

 1

 10

 1

 8

 1

 8

 1

 5

 1

 4

 1

 3

 1

 1

 1

 1

 1

 1

 1

 3

 1

 2

 1

 1

 1

 1

 1

 1

 1

 1

 1

 2

 1

 2

 1

 2

 1

 2

 1

 4

 1

 4

 1

 4

 1

 4

 1

 4

 1

 316

 1

 316

 1

 286

 1

 188

 1

 188

 1

 188

 1

 188

 1

 53

 1

 2

 1

 35

 1

 5

 1

 12

 1

 4

 1

 36

 1

 11

 1

 5

 1

 17

 1

 4

 1

 2

 1

 2

 1

 98

 1

 7

 1

 7

 1

 7

 1

 7

 1

 83

 1

 83

 1

 1

 1

 1

 1

 1

 1

 1

 1

 56

 1

 22

 1

 8

 1

 3

 1

 1

 1

 4

 1

 2

 1

 2

 1

 1

 1

 1

 1

 27

 1

 26

 1

 1

 1

 4

 1

 3

 1

 2

 1

 1

 1

 1

 1

 1

 1

 1

 1

 1

 1

 1

 1

 1

 1

 1

 1

 1

 1

 7

 1

 3

 1

 1

 1

 1

 1

 1

 1

 1

 1

 12

 1

 5

 1

 1

 1

 1

 1

 1

 1

 2

 1

 1

 1

 1

 1

 6

 1

 5

 1

 1

 1

 8

 1

 8

 1

 8

 1

 3

 1

 5

 1

 28

 1

 28

 1

 7

 1

 7

 1

 7

 1

 2

 1

 3

 1

 2

 1

 20

 1

 3

 1

 2

 1

 2

 1

 1

 1

 1

 1

 1

 1

 1

 1

 1

 1

 16

 1

 16

 1

 3

 1

 7

 1

 2

 1

 1

 1

 1

 1

 2

 1

 1

 1

 1

 1

 1

 1

 1

 1

 2

 1

 2

 1

 2

 1

 2

 1

 2

 1

 2

 1

 220

 1

 133

 1

 27

 1

 27

 1

 8

 1

 8

 1

 8

 1

 19

 1

 19

 1

 19

 1

 104

 1

 37

 1

 37

 1

 30

 1

 2

 1

 3

 1

 2

 1

 14

 1

 14

 1

 7

 1

 7

 1

 52

 1

 52

 1

 52

 1

 1

 1

 1

 1

 2

 1

 2

 1

 2

 1

 1

 1

 1

 1

 1

 1

 1

 1

 1

 1

 1

 1

 69

 1

 5

 1

 5

 1

 2

 1

 2

 1

 3

 1

 3

 1

 64

 1

 53

 1

 45

 1

 10

 1

 9

 1

 9

 1

 9

 1

 1

 1

 1

 1

 30

 1

 30

 1

 30

 1

 5

 1

 5

 1

 5

 1

 5

 1

 8

 1

 3

 1

 3

 1

 3

 1

 3

 1

 5

 1

 5

 1

 3

 1

 3

 1

 3

 1

 2

 1

 2

 1

 11

 1

 11

 1

 11

 1

 6

 1

 6

 1

 5

 1

 5

 1

 18

 1

 18

 1

 18

 1

 18

 1

 18

 1

 18

 1

 45

 1

 32

 1

 28

 1

 11

 1

 9

 1

 9

 1

 8

 1

 8

 1

 4

 1

 4

 1

 8

 1

 3

 1

 1

 1

 4

 1

 5

 1

 5

 1

 1

 1

 1

 1

 1

 1

 1

 1

 1

 1

 1

 1

 9

 1

 9

 1

 9

 1

 9

 1

 9

 1

 9

 1

 1

 1

 8

 1

 7

 1

 1

 1

 22

 1

 22

 1

 3

 1

 3

 1

 3

 1

 3

 1

 19

 1

 1

 1

 1

 1

 1

 1

 1

 1

 1

 1

 1

 1

 18

 1

 18

 1

 14

 1

 14

 1

 14

 1

 4

 1

 1

 1

 1

 1

 3

 1

 3

 1

 1

 1

 1

 1

 1

 1

 1

 1

 1

 1

 1

 1

 1

 1

 1

 1

 1

 1

 1

 1

 1

 1

 1

 1

 2878

 1

 2734

 1

 1424

 1

 693

 1

 342

 1

 288

 1

 7

 1

 1

 1

 20

 1

 3

 1

 4

 1

 14

 1

 9

 1

 23

 1

 45

 1

 4

 1

 5

 1

 8

 1

 4

 1

 4

 1

 1

 1

 3

 1

 3

 1

 6

 1

 92

 1

 78

 1

 8

 1

 3

 1

 2

 1

 228

 1

 185

 1

 11

 1

 7

 1

 1

 1

 1

 1

 2

 1

 2

 1

 1

 1

 3

 1

 1

 1

 2

 1

 52

 1

 17

 1

 8

 1

 6

 1

 2

 1

 2

 1

 2

 1

 4

 1

 4

 1

 4

 1

 4

 1

 27

 1

 55

 1

 36

 1

 36

 1

 18

 1

 4

 1

 3

 1

 13

 1

 2

 1

 15

 1

 15

 1

 15

 1

 15

 1

 835

 1

 74

 1

 22

 1

 1

 1

 1

 1

 478

 1

 455

 1

 154

 1

 52

 1

 9

 1

 11

 1

 27

 1

 6

 1

 1

 1

 19

 1

 1

 1

 194

 1

 4

 1

 4

 1

 6

 1

 3

 1

 3

 1

 55

 1

 55

 1

 54

 1

 19

 1

 101

 1

 92

 1

 209

 1

 58

 1

 35

 1

 1

 1

 1

 1

 132

 1

 28

 1

 28

 1

 14

 1

 10

 1

 1

 1

 17

 1

 4

 1

 4

 1

 4

 1

 7

 1

 4

 1

 87

 1

 87

 1

 2

 1

 36

 1

 22

 1

 3

 1

 2

 1

 11

 1

 30

 1

 12

 1

 3

 1

 5

 1

 2

 1

 1

 1

 11

 1

 1

 1

 1

 1

 1

 1

 1

 1

 5

 1

 4

 1

 2

 1

 2

 1

 1

 1

 3

 1

 3

 1

 1

 1

 1

 1

 1

 1

 1

 1

 4

 1

 1

 1

 1

 1

 1

 1

 6

 1

 2

 1

 1

 1

 1

 1

 1

 1

 3

 1

 3

 1

 17

 1

 6

 1

 1

 1

 1

 1

 1

 1

 3

 1

 2

 1

 2

 1

 1

 1

 1

 1

 5

 1

 2

 1

 2

 1

 1

 1

 1

 1

 1

 1

 1

 1

 1

 1

 1

 1

 1

 1

 5

 1

 3

 1

 1

 1

 2

 1

 2

 1

 2

 1

 2

 1

 2

 1

 1

 1

 1

 1

 1

 1

 1

 1

 1

 1

 25

 1

 9

 1

 9

 1

 9

 1

 9

 1

 9

 1

 1

 1

 3

 1

 5

 1

 1

 1

 1

 1

 1

 1

 1

 1

 1

 1

 1

 1

 1

 1

 5

 1

 2

 1

 2

 1
